# Exodermis lignification impacts lateral root emergence in *Brachypodium distachyon*

**DOI:** 10.1101/2025.09.02.673693

**Authors:** Kevin Bellande, Cristovāo De Jesus Vieira Teixeira, Angelina D’orlando, Léa Perez, Richard Sibout, Anne Roulin, Joop E.M. Vermeer, Thomas Badet

## Abstract

**Rationale:** The mechanisms controlling lateral root emergence in monocots, particularly the role of the exodermis, are poorly understood. We investigated how natural variation in the *Brachypodium distachyon* stress response shapes root system architecture by modulating cell wall dynamics.

**Methods:** We used root tip excision to synchronize lateral root development across natural accessions. The resulting phenotypes were analysed using comparative transcriptomics, biochemical lignin quantification, confocal Raman spectroscopy, and chemical inhibition of lignin biosynthesis.

**Key Results:** Two distinct root system architectures, ’pine tree’ and ’fishbone’, were identified. The ’fishbone’ phenotype results from an emergence-specific defect caused by the premature and intense lignification of the exodermis. This was driven by the transcriptional upregulation of lignin biosynthesis genes and was rescued by a lignin inhibitor.

**Main Conclusion:** Stress-induced exodermal lignification acts as a mechanical ’brake’ on lateral root emergence. This positions the exodermis as a key regulatory hub that integrates environmental cues with developmental programs to control RSA plasticity in grasses.

## Introduction

Plants exhibit remarkable intraspecific phenotypic variation, enabling individuals within a species to respond and adapt to diverse environmental conditions. In plants, natural variation has been extensively characterized for above-ground traits such as flowering time and abiotic stresses (Filiz *et al*., 2009; Schwartz *et al*., 2010; Luo *et al*., 2011; Ingram *et al*., 2012; González *et al*., 2020; Stritt *et al*., 2022; Ludwig *et al*., 2024). Increasing evidence also points to substantial variation in below-ground traits, particularly root system architecture (Chochois *et al*., 2015; Rouina *et al*., 2025). Indeed, root system architecture plasticity is a key component of plant adaptation to changing environmental conditions, enabling dynamic modulation of water and nutrient foraging strategies (Lynch et al., 1995; Smith & De Smet, 2012; Maqbool et al., 2022). A central mechanism driving root plasticity is the formation of lateral roots, which remodel the root system in response to developmental and environmental cues (Lynch, 2019; Ramachandran *et al*., 2024).

Lateral root development requires coordinated cellular processes such as cell division, expansion, and differentiation, that occur within the context of a mechanically complex tissue (Beckers *et al*., 2024). To emerge, lateral root must traverse several established cell layers of the primary root, posing significant physical constraints (Stoeckle *et al*., 2018; Vadodaria & Anderson, 2025). These constraints are mediated by plant cell walls, which are composed of polysaccharide polymers such as cellulose, hemicellulose, and pectin (in dicot primary cell walls), and often reinforced with lignin, particularly in secondary cell walls of differentiated tissues like the endodermis and exodermis (Sarkar *et al*., 2009; Geldner, 2013; Voxeur *et al*., 2015; Höfte & Voxeur, 2017; S Artur *et al*., 2021; Liu & Kreszies, 2023). While these cell layers serve as structural barriers and diffusion regulators, they must also undergo regulated remodelling to allow lateral root emergence (Shang *et al*., 2025; Vadodaria & Anderson, 2025).

In *Arabidopsis thaliana*, the molecular pathways controlling cell wall remodelling during lateral root initiation from xylem pole pericycle cells are well established (Swarup *et al*., 2008; Péret *et al*., 2012; Kumpf *et al*., 2013; Lucas *et al*., 2013; Lewis *et al*., 2013; Vermeer *et al*., 2014; Roycewicz & Malamy, 2014; Berhin *et al*., 2019; Wachsman *et al*., 2020; Ursache *et al*., 2021). However, the regulation and evolutionary variability of these mechanisms in monocots, which possess distinct root anatomies, remain poorly understood (Petrova *et al*., 2023; Zhu *et al*., 2025). In *Brachypodium distachyon*, lateral root formation involves not only the pericycle but also contributions from the endodermis and cortex (De Jesus Vieira Teixeira et al., 2024). To emerge, lateral root must ultimately overcome the exodermis, a suberin- and lignin-rich outer barrier whose role during organogenesis remains unclear (Liu & Kreszies, 2023). The composition and mechanical properties of the exodermis, particularly its lignification patterns, vary across species and are likely to influence lateral root development (Voxeur *et al*., 2015; Cantó-Pastor *et al*., 2024; Manzano *et al*., 2024).

Here, we propose that the exodermis functions not only as a structural constraint but also as a regulatory interface that integrates stress signals to modulate root development. We use root tip excision to synchronize lateral root development and uncover natural variation in root system architecture between two *B. distachyon* accessions, Bd21 and Bd21-3. Through comparative transcriptomics, cell wall profiling, and microscopy, we demonstrate that these phenotypic differences are driven by distinct cell wall remodelling processes, with differential lignification of the exodermis emerging as a primary determinant of emergence.

## Materials and Methods

### Plant materials and growth conditions

A total of 23 *B. distachyon* accessions were used in this study: Bd21, Bd21-3, SAP47, CM3, BdtR7a, TEK2, GES1, BdTR8C, ko21, Adi2, BdTRIIA, BdTRIIG, BdTR10C, Fo21, BdTR9K, GAZ8, ABR7, PER4, Lam13, Bd30-1, RON2, CRO24, and SAN11, as previously described (Stritt *et al*., 2022). Seeds were dehulled and surface sterilized in 6% (v/v) bleach containing 0.01% Triton™ X-100 (Sigma-Aldrich, Ref: X100-5ML) for 1 minute, then rinsed six times with sterile water. Sterilized seeds were sown on square 12 × 12 cm Petri dishes containing 1% plant agar (w/v) supplemented with half-strength Murashige and Skoog (½ MS) basal medium and buffered with MES. No more than 15 seeds were placed in a single row approximately 3.5 cm from the top edge of each plate, with embryos facing upward and avoiding direct contact with the medium surface. Plates were placed vertically at an angle of ∼20° in growth chambers under continuous light at 22°C to promote root growth along the surface. For root tip excision experiments, 6-day-old seedlings had approximately 0.5 cm of the primary root tip removed using a sterile scalpel. After excision, plates were resealed and returned to the same growth conditions to allow for synchronized lateral root formation. Samples for RNA sequencing were collected at 0, 1, 2, 4, 8, and 12 hours following root tip excision. For each time point, a 5 mm segment of the root, immediately proximal to the cut, was harvested from a pool of 20-30 seedlings. All samples were immediately flash-frozen in liquid nitrogen upon collection. For chemical inhibition of lignin biosynthesis, 6-day-old seedlings were grown on ½ MS 1% agar plates. After the root tip excision, a sterile filter paper saturated with a 10 µM solution of piperonylic acid was gently applied to the whole root system. Treated seedlings of both Bd21 and Bd21-3 accessions were maintained under growth chamber conditions for 60 hours prior to phenotypic observation and quantification of lateral root numbers.

### RNA extraction, library preparation, and sequencing

For the *B. distachyon* Bd21 and Bd21-3 accessions, total RNA was extracted from root tips after excision using the SV Total RNA Isolation System (Promega) according to the manufacturer’s instructions. The RNA samples were shipped to Novogene (NOVOGENE (UK) company limited) for quality control, library preparation, and high-throughput sequencing. RNA integrity and purity were first assessed by Novogene’s in-house quality control pipeline, and only samples with sufficient RNA integrity number and concentration passed to library preparation. Messenger RNA was enriched from total RNA using poly-T oligo-attached magnetic beads, followed by fragmentation and first-strand cDNA synthesis with random hexamer primers. Second-strand cDNA synthesis, end-repair, A-tailing, adapter ligation, and PCR enrichment were performed to generate sequencing-ready libraries. Libraries were prepared as unstranded, paired-end libraries and sequenced on an Illumina NovaSeq 6000 platform, yielding 150 base-pair paired-end reads (PE150). Approximately 6 Gb of raw sequencing data per sample were generated, with >85% of bases expected to reach Q30 quality standards.The generated raw reads were first filtered for polyA and TruSeq adapters using the BBduck tool from the BBMap suite (v38.96) (options ref=polyA.fa.gz,truseq.fa.gz ktrim=r k=23 mink=11 hdist=1 tpe tbo qtrim=r trimq=10 minlength=20) (Bushnell *et al*., 2014). The resulting trimmed reads were next aligned to the respective genome assembly (BdistachyonBd21_3_537_v1.0 for Bd21-3, and Bdistachyon_314_v3 for Bd21) using STAR aligner (v2.7.10a) (Dobin *et al*., 2013). The genome index was built using strings of 12 bases (--genomeSAindexNbases 12), and reads were aligned allowing for multiple matches (options -- outFilterMultimapNmax 100 --winAnchorMultimapNmax 200) (Li *et al*., 2009). The coordinate-sorted BAM files were indexed with samtools (v1.15.1), and read counts were calculated using the union method implemented by the htseq-count tool from the HTSeq suite (v2.0.2) using version 1.2 of the Bd21-3 genome annotation (BdistachyonBd21_3_537_v1.2.gene.gff3) (Anders *et al*., 2015).

### Pangenome reconstruction

To identify homologs between the annotated genes in Bd21 and Bd21-3 accessions, we first extracted all coding sequences given the gene annotations using *gffread* v0.12.7 (BdistachyonBd21_3_537_v1.2 and Bdistachyon_556_v3.2 available at the Phytozome version 14 database) (Pertea & Pertea, 2020). The transcripts were translated into proteins using the *transeq* tool from the EMBOSS:6.6.0.0 suite and orthology between Bd21 and Bd21-3 proteins was finally inferred using *OrthoFinder* version 2.5.5 with default parameters (-og -a 12) (Rice *et al*., 2000; Emms & Kelly, 2019).

### Differential expression analysis

Differential expression analysis was performed using R version 4.1.2 and the DESeq2 package version 1.32.0 (‘R: The R Project for Statistical Computing’; Love *et al*., 2014). Genes with null expression in more than 20% of the samples were filtered out. The data dispersion was estimated using the *vst* function implemented in DESeq2 package using a sample of 20,000 genes (nsub=20000). Differential expression across time-points was calculated on genes expressed in at least 20% of the samples and with a total read count >= 10. We used the likelihood ratio test to estimate changes in gene expression across the kinetic and extracted results for each comparison using the Wald test implemented in DESeq2.

### Gene ontology enrichments

All gene ontology (GO) enrichment analyses were performed using the R package *topGO* version 2.44.0 based on the Bd21 and Bd21-3 annotations (Bdistachyon_556_v3.2.annotation_info.txt and BdistachyonBd21_3_537_v1.2.annotation_info.txt available on Phytozome version 14 database. Results are shown at the biological processes (BP) level at a false discovery rate of 5% for GOs annotated in >10 genes and with >5 occurrences in the test gene set. Enrichment values are expressed as the number of occurrences of the GO in the test gene set over its Fisher test expected number.

### Preparation of hand-sectioned root samples

For sectioning, roots from 6-old seedlings of similar length were placed in parallel. Fragments of 1 cm containing the region of interest were partitioned and embedded in 4% agarose. Once solidified, the agarose blocks were glued onto a hand microtome, and sections of approximately 50 µm were prepared for clearing or immediate visualization. Representative images were obtained from at least 30 seedlings from three independent replicates.

### Clearing and staining

Clearing steps using DEEP-Clear were performed as described in (Pende et al., 2020; De Jesus Vieira Teixeira et al., 2024). The DEEP-Clear solution consists of 5% to 8% (v/v) THEED, 5% (v/v) Triton X-100, and 25% (w/v) urea in water. Seven-day-old root seedlings were collected and fixed for 1 hour in 4% (w/v) paraformaldehyde in 1× phosphate-buffered saline (PBS) with three rounds of soft vacuum infiltration. After fixation, roots were washed five times in 1× PBS. Samples were then transferred to the DEEP-Clear solution and incubated at room temperature with gentle shaking for 7-10 days, with the solution being replaced twice. For staining, stock solutions (1%) of Basic Fuchsin (lignin), Renaissance, and/or Calcofluor (cell wall) were prepared directly in DEEP-Clear. To combine multiple dyes, samples were incubated first in Basic Fuchsin (0.1% in DEEP-Clear) for 1 hour and washed overnight. Samples were then transferred to Renaissance (0.1% in DEEP-Clear) for 2 days. To visualize lignified tissues, *B. distachyon* root segments were stained with freshly prepared Wiesner stain (phloroglucinol-HCl). The staining solution was made by dissolving 0.3 g of phloroglucinol in 10 ml of 100% ethanol and then mixing this solution with concentrated HCl at a 2:1 (v/v) ratio. Root sections were incubated in the stain for approximately 10 minutes at room temperature. The stained sections were immediately mounted on a microscope slide in a drop of the staining solution and observed with a light microscope. Due to the transient nature of the stain, images were captured within 10 minutes of application to prevent signal deterioration.

### Quantification of lignin

For the CASA lignin method, three independent replicates were prepared for Bd21 and for Bd21-3. Thirty whole roots were collected (10 seedlings per replicate in three square dishes), pooled, frozen in liquid nitrogen, and ground for 2 minutes at full speed. The samples were then dried in 2 mL Eppendorf tubes at 40°C for 24 hours. A cysteine stock solution (0.1 g mLLJ¹) in 72% sulfuric acid (SA) was prepared by dissolving 10 g L-cysteine in 100 mL SA. The solvent-extracted biomass sample (∼5 mg) was placed in a 4 mL glass vial, and 1.0 mL of the cysteine stock solution was added. The mixture was sealed with a Teflon-lined screw cap and stirred at 24°C (500 rpm) for 60 min until complete biomass dissolution. The solution was then diluted 1/50 by adding 20 µL of the solution to 980 µL of deionized water in a 4 mL vial. The absorbance at 283 nm (ALJLJLJ) was measured using a UV spectrophotometer against a blank. If necessary, a UV absorption spectrum was recorded from 230 to 400 nm at 1 nm intervals. Lignin content (CASA_L%) was calculated using the Beer–Lambert law. All tests were performed in triplicate, and data are presented as mean ± SD.

For the thioacidolysis, root segments from 30–50 individual roots were collected from two distinct regions: a proximal region (5 mm above the apex after root tip excision) and an upper region (from 5 mm above the apex to the root collar). Samples were pooled, weighed, and lyophilized. The thioacidolysis reagent was prepared by mixing 2.5 mL BFLJ etherate (Aldrich) and 10 mL ethane thiol (EtSH, Aldrich) in a 100 mL flask, adjusting the final volume to 100 mL with dioxane (pestipur grade). The extract-free biomass (∼5 mg) was incubated with 3 mL of reagent and 0.1 mL of an internal standard solution (2.5 mg mLLJ¹ heneicosane in CHLJClLJ) in a Teflon-lined screw-capped glass tube. Thioacidolysis was conducted at 100°C for 4 h. After cooling, the pH of the reaction mixture was adjusted using 0.2 M NaHCOLJ, followed by the addition of 0.1 mL of 6M HCl. The mixture was extracted with 2 mL of CHLJClLJ, and after phase separation, the lower organic phase was recovered and dried over NaLJSOLJ. The solvent was evaporated under reduced pressure at 40°C until approximately 0.5 cm of liquid remained in the vial before silylation for GC or GC-MS analysis. Lignin-derived dimers were quantified after desulfurization, and monomer analysis was performed as described (Lapierre & Rolando, 1988).

### Confocal Raman spectroscopy

For the sample preparation and spectral acquisition, reference peaks for major cell wall components (e.g., cellulose, hemicellulose, and lignin) were identified by comparison to established literature spectra (Diehn *et al*., 2024). For the analysis, seven-day-old *B. distachyon* seedlings (accessions Bd21 and Bd21-3) were grown in vitro and sampled at 0 h and 40 h after root tip excision. Root segments from phenotypically distinct upper root regions were excised, aligned, and embedded in 8% agarose. Transverse sections of 50 µm were obtained using a vibratome (HM 650 V, Microm Microtech), air-dried for 10 min, and mounted on Raman slides. Spectra were acquired using a confocal Raman microscope (inVia Reflex, Renishaw) with a 785 nm laser. The system was equipped with a double-edge filter (Rayleigh cutoff at 100 cmLJ¹) and two diffraction gratings (1800 and 1200 L/mm) for spectral acquisition. An automated XYZ stage allowed for precise 100 nm step displacements. Laser power was optimized to prevent sample damage. Spectra were collected with a resolution of <4 cmLJ¹ and a precision of <0.5 cmLJ¹. Measurements were focused on the anticlinal cell walls of the exodermis, with at least 15 root sections analyzed from five independent roots per genotype, across three to five biological replicates. Raw spectral data quality was first enhanced by applying cosmic ray removal using Wire 4.2 software (Renishaw). Subsequent pre-processing was conducted in OriginLab Pro, which included baseline correction and normalization of the spectra to ensure comparability. Following these steps, an average Raman spectrum was generated for each sample for further analysis. All downstream statistical analyses and data visualization were performed in R (v4.4.2) utilizing the tidyverse suite of packages. To handle technical duplicates, spectra from replicate measurements for each sample were averaged at each wavenumber. To ensure comparability, only spectra covering the full range from 300 to 1800 cmLJ¹ were included in the final analysis. To identify statistically significant differences in cell wall composition between the Bd21 and Bd21-3 accessions, the mean intensity of major cell wall-related peaks was compared for each tissue type using an independent two-sample t-test. A difference was considered statistically significant at *p* < 0.05.

## Results

### Differential lateral root emergence dynamics in response to primary root tip excision

To explore the natural variation in lateral root formation in *B. distachyon*, we employed a root tip excision assay, a method previously shown to synchronize lateral root formation (Pacheco-Villalobos *et al*., 2013). Focusing on a set of 23 natural accessions spanning the species distribution along the Mediterranean range, the approach revealed two distinct root system architecture phenotypes. The reference accession Bd21 together with 7 other accessions displayed a “pine tree” phenotype, characterized by uniformly spaced lateral roots along the primary root axis four days after excision (BdTR8i, Tek-2, Sap47, Ges1, BdTR11A, BdTR11G and Gaz-8). In contrast, Bd21-3 and 14 other accessions exhibited a “fishbone” phenotype, with lateral roots clustered near the excision site but largely absent in the more upper regions of the primary root (Figs.LJ1a, S1). The two phenotypes segregated in the A-Italia, A-East and B-East genetic and geographical clusters while accessions from the B-West and C-Italia clusters all displayed the “fishbone” phenotype, suggesting that it might underlie ecological adaptation (Fig.LJ1b) (Stritt *et al*., 2022).

To determine whether the “fishbone” or the “pine tree” phenotypes reflected a constitutive developmental program or a specific response to root tip excision, we first compared the root system architecture of two representative accessions under control (non-excised) conditions. Non-excised 6d-old Bd21 accession (pine tree) seedlings developed significantly longer primary roots and a higher number of total lateral roots than Bd21-3 (fishbone) (Fig.LJ1c). However, lateral root density was similar between the two accessions, indicating that the greater number of lateral roots in Bd21 is likely due to its longer primary roots rather than a higher branching frequency (Fig.LJ1c). We next analysed the spatial distribution of emerged lateral roots along the primary root axis of Bd21 and Bd21-3 plants after root tip excision. In both accessions, lateral roots were initiated close to the excision site, and within the 1^st^ centimetre of the root axis their development appeared comparable. However, beyond 2 cm from the excision point, Bd21-3 exhibited markedly fewer emerged lateral roots. In Bd21, primary root tip excision resulted in consistent lateral root emergence along the root axis, with density gradually declining only near the root– shoot junction. By contrast, in Bd21-3 at 72 hours after root tip excision, lateral roots in the upper part of the primary root either failed to emerge or failed to grow into the nutrient medium (Fig. 1d). To gain further insights into putative defects in lateral root emergence, we followed the temporal progression of lateral root development after the root tip excision over a 32-hour time course (1 mm, 2 mm, and 3LJmm above the excision site) (De Jesus Vieira Teixeira et al., 2024). Both Bd21 and Bd21-3 displayed synchronized lateral root development, with early stages detectable 2LJhours post excision (stage 1-2) and nearly all primordia reaching late developmental stages by 24 h (stage 9-10) (Fig.LJS2).

**Fig. 1.**
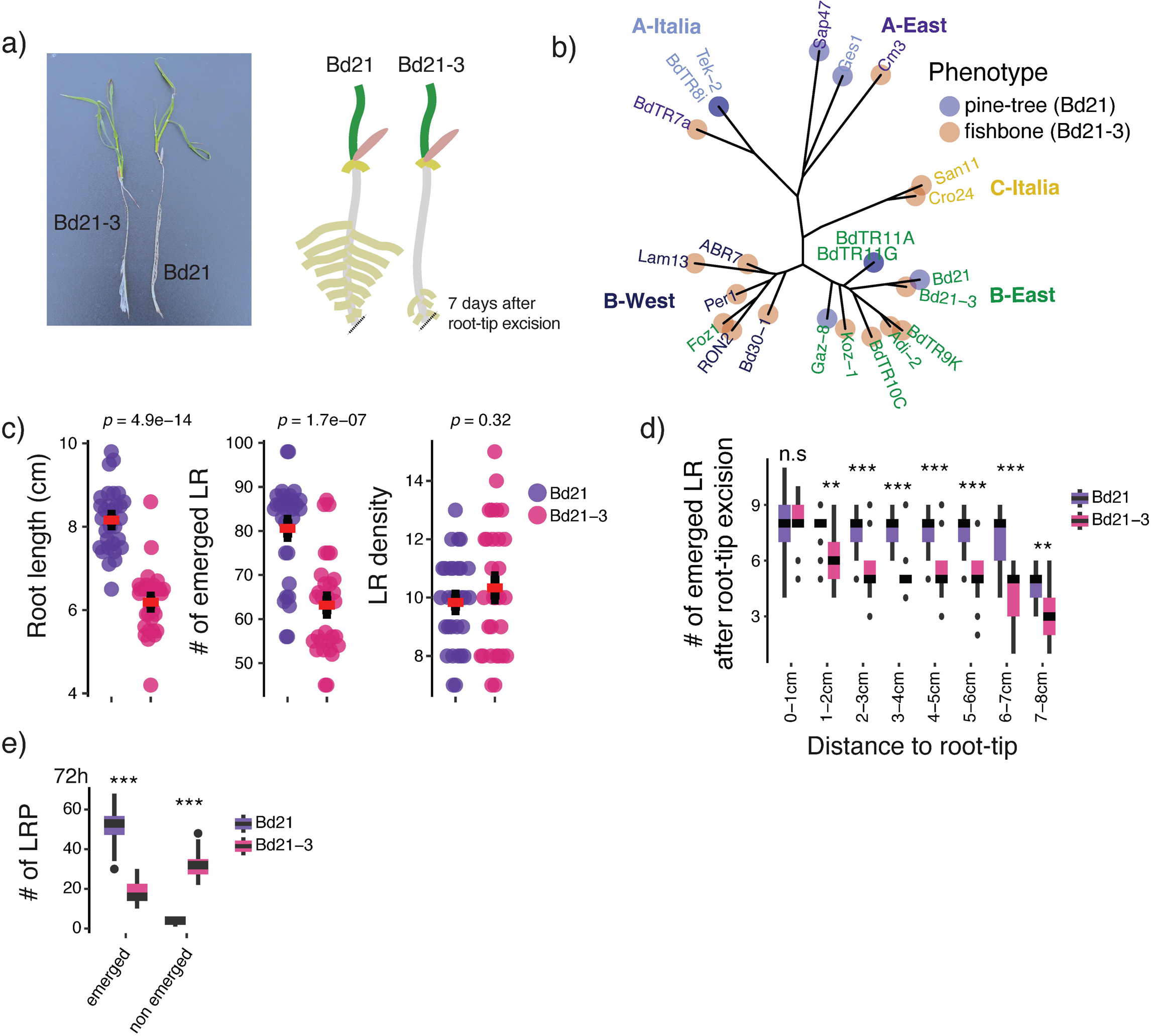
Bd21 and Bd21-3 display distinct lateral root emergence dynamics following root tip excision. (a) Representative images of *B.distachyon* accessions Bd21 and Bd21-3 phenotypes and related schematic, 4 days after root tip excision. (b) Geographically structured distribution of the “pine tree” and “fishbone” root system architecture phenotypes across a panel of 23 natural *B.distachyon* accessions, indicated geographical clusters (colour codes on the tree). (c) Quantification of root system architecture traits under non-excised conditions. (d) Spatial distribution of emerged lateral roots along the primary root axis after root tip excision. (e) Quantification of emerged versus non-emerged lateral root primordia in the root of Bd21 and Bd21-3. * Error bars represent standard deviation. Asterisks indicate statistical significance based on a Student’s t-test (p < 0.05, *p < 0.01, *p < 0.001).

To directly address whether the “fishbone” phenotype in Bd21-3 results from a defect in lateral root emergence, we quantified the number of emerged and non-emerged lateral root 72 hours after root tip excision in the primary root of both accessions. We find that Bd21-3 exhibited a significantly higher proportion of non-emerged lateral root in the upper region compared to Bd21 (Figs.LJ1e, S3). These results indicate that lateral root initiation and development are not impaired in Bd21-3, confirming that the primary defect lies in lateral root emergence. Results on non-excised roots further support the notion that the ’fishbone’ phenotype observed in Bd21-3 is not intrinsic but rather arises from a specific defect in lateral root emergence that is triggered or exacerbated by root tip excision.

### Comparative transcriptomic analysis reveals extensive cell wall remodelling following root tip excision

To investigate the genetic basis underlying the difference between the “pine tree” and the “fishbone” lateral root phenotypes, we first performed a comparative genomic analysis using Bd21 and Bd21-3 fully annotated genomes. We assigned the full set of 103,438 proteins to 38,403 ortholog groups (orthogroups), from which ∼73% are single-copy orthogroups present in both accessions (28,276 / 38,403, Fig. 2a). The Bd21-3 accession has 3,963 orthogroups for which we could not find an ortholog in Bd21. Conversely, 1,814 orthogroups are exclusively found in the Bd21 accession. A total of 3,138 orthogroups have a single protein representative in one of the two accessions while the other has additional proteins attached to the same orthogroup, suggesting recent duplication events in both accessions (Fig. 2a). Orthogroups for which multiple proteins are found in Bd21 but only one in Bd21-3 are enriched for functions related to DNA repair and stress response and might contribute to the phenotypic difference we observe after primary root tip excision (Fig. S4). Thus, the two-accession pangenome reveals substantial gene content variation in *B. distachyon*.

**Fig. 2.**
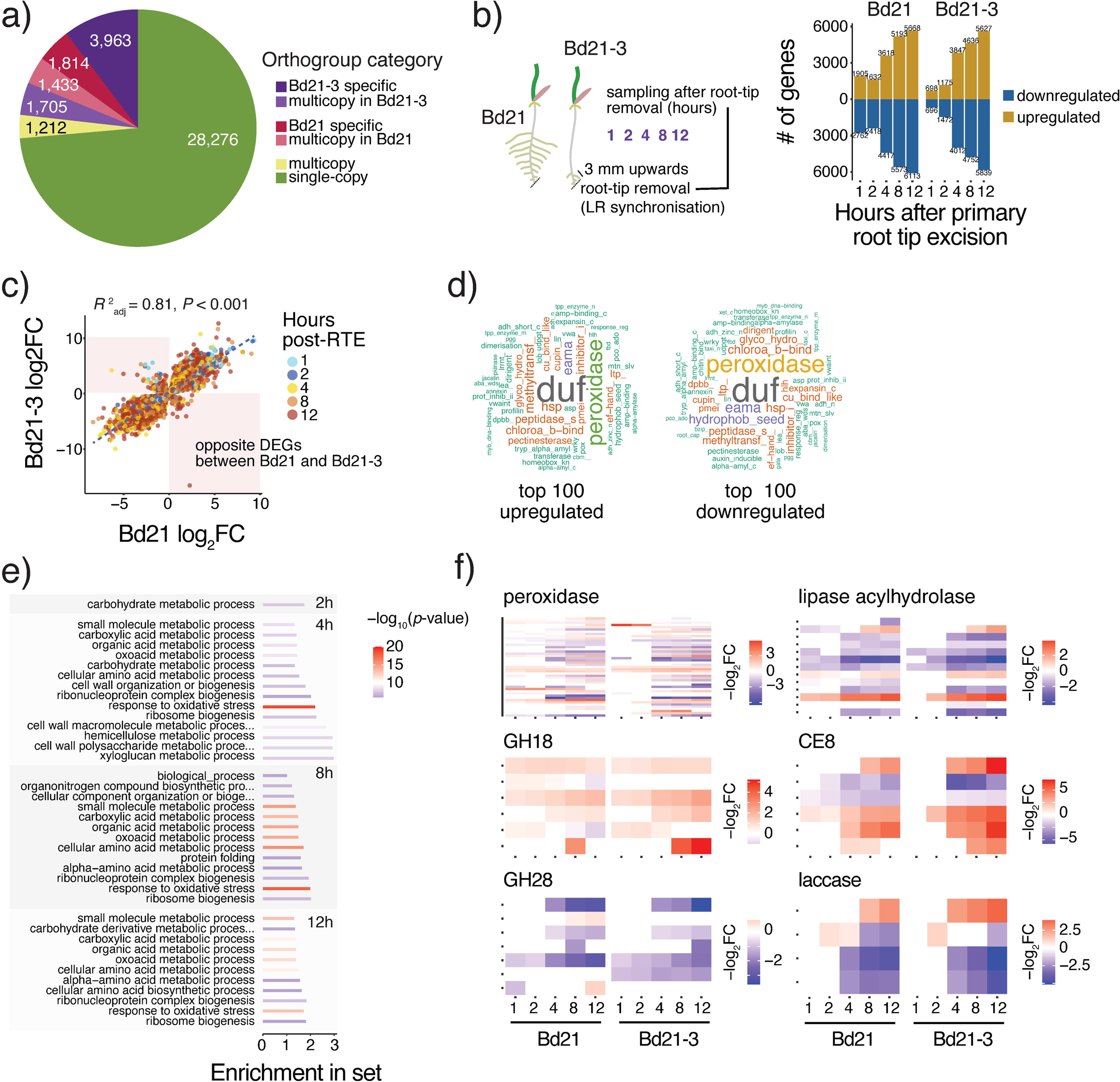
Comparative genomic and transcriptomic analysis reveals a divergent cell wall remodelling response in Bd21 and Bd21-3 after root tip excision. (a) Pie chart showing the pangenome composition of Bd21 and Bd21-3, with colours indicating the distribution of orthogroup categories. (b) Schematic representation of the time-course transcriptomic experiment following root tip excision for lateral root primordia synchronization (left panel), and the corresponding number of significantly up- and downregulated genes at 1, 2, 4, 8, and 12 hours post excision in both accessions (right panel). (c) Correlation plot of logLJ fold-change (logLJFC) values for differentially expressed single-copy orthologs between Bd21 and Bd21-3. Points are coloured by time point post excision, with a subset of genes showing opposite regulation highlighted in red. (d) Word clouds representing the most frequent annotation terms among the top 100 upregulated and top 100 downregulated genes. (e) Gene Ontology (GO) enrichment analysis of similarly differentially expressed genes at 2, 4, 8, and 12 hours after root tip excision in both accessions. The colour scale indicates (–logLJLJ(*p*-value)) of enrichment. (f) Heatmaps showing the fold-change value of differentially expressed genes over time for selected cell wall–related enzyme families in both accessions (–log_2_FC).

To identify the cellular processes involved in the “pine tree” and “fishbone” lateral root phenotypes, we next performed a time-course transcriptomic analysis of the root tissue (at 0h; 1h; 2h; 4h; 8h; 12h) after root tip excision in Bd21 and Bd21-3 accessions (Fig. 2b). Approximately 44% and 35% of the annotated genes in Bd21 and Bd21-3 respectively, showed expression in the root tissue in at least one time point (with >10 read counts). Principal component analysis revealed two major transcriptomic clusters corresponding to early (0 to 2 hours) and late time points (4 to 12 hours) after synchronization (Fig. S5). Compared to time 0 after synchronization, differential gene expression became more pronounced at the later time points. The number of up- and downregulated genes progressively increased over time, reaching between 3,618 and 6,113 differentially expressed genes (DEGs) between 4 and 12 hours after synchronization (Fig. 2b). At all-time points, Bd21 exhibited more DEGs than Bd21-3, likely reflecting the higher sequencing depth in Bd21 samples.

Focusing on 7,980 single-copy orthogroups differentially expressed in both accessions, we found that fold-change values were strongly correlated between Bd21-3 and Bd21, suggesting a largely conserved transcriptional response to root tip excision (Fig. 2c, R^2^ = 0.81, p-value < 0.001). Gene Ontology (GO) enrichment analysis of differentially expressed genes in both accessions revealed a strong overrepresentation of terms related to oxidoreductase and peroxidase activity (Fig. S6). In addition, we identified a set of 240 orthologs that exhibited inverse expression changes between Bd21 and Bd21-3 over the time course (Fig. 2c). GO enrichment analysis of these divergently regulated genes showed a significant enrichment for energy-intensive processes, such as ATP hydrolysis (Fig. S6). Further analysis of the top 100 most upregulated and downregulated genes in both accessions revealed consistent enrichment for cell wall-related enzymes, including glycosyl hydrolases, pectinesterases, and expansins. Notably, peroxidases emerged as the most prominently enriched category (Fig. 2d). In agreement, GO terms associated with carbohydrate metabolism and cell wall biogenesis were also significantly enriched at genes similarly differentially expressed in both accessions at early time points after root tip excision (Fig. 2e).

To further explore the contribution of cell wall related metabolism to lateral root emergence, we focused on genes encoding for proteins listed in the plant cell wall experimental database (Figs. 2f, S7). Approximately ∼87% of the expressed genes mapped to this database were differentially expressed after root tip excision (484 out of 555 genes). Among the most deregulated protein families were peroxidases and proteins of unknown function. Several glycoside hydrolases (GHs) were also significantly mis-regulated following synchronisation. Notably, 92% of the differentially expressed GH family 18 genes, which include chitinases, were upregulated after root tip excision (37 out of 40). In contrast, 90% of the GH family 28 genes, which encode polygalacturonases involved in pectin remodelling, were downregulated together with many arabinogalactan proteins (30 out of 33; Figs. 2f, S8). In contrast, pectin methylesterase and pectin methyltransferase inhibitors (61.5% and 90.9% up-regulated respectively), expansin (70.5% up-regulated), and glucan-acting glycosyl hydrolases (GH17, 77.5% up-regulated), were predominantly upregulated. The differential regulation of these types of cell wall modifying enzymes in Bd21-3 might underly increased rigidity of the cell wall in the root tissue. Altogether, the transcriptomic analysis reveals that root tip excision triggers extensive cell wall remodelling in both *B. distachyon* accessions, with differences in timing, magnitude, and specific gene families likely underlying the difference between the “pine tree” and the “fishbone” phenotypes of lateral root emergence.

### Differential lignification in the exodermis underlies the “fishbone” root system architecture

To investigate potential differences in cell wall modifications following lateral root synchronization, we employed a histochemical approach to visualize major cell wall polymers. Given the strong enrichment of peroxidase-related GO terms in our transcriptomic data, we specifically examined lignin accumulation using phloroglucinol-HCl staining (Figs. 3a, S9). At 40 hours post excision, Bd21-3 roots exhibited stronger lignin staining along the root axis and vasculature compared to Bd21, suggesting greater overall lignin accumulation. To biochemically validate the visual increase in lignin content in Bd21-3 roots, we quantified total lignin content using the Cysteine-Assisted Sulfuric Acid (CASA) method. This revealed no significant difference in control lignin levels (0h) between the two accessions (Figs. 3b, S10). However, 40 hours after root tip excision, Bd21-3 roots exhibited a significant increase in lignin content (18.71 ± 0.94%) compared to Bd21 (15.33 ± 0.50%, *p* = 0.0201).

**Fig. 3.**
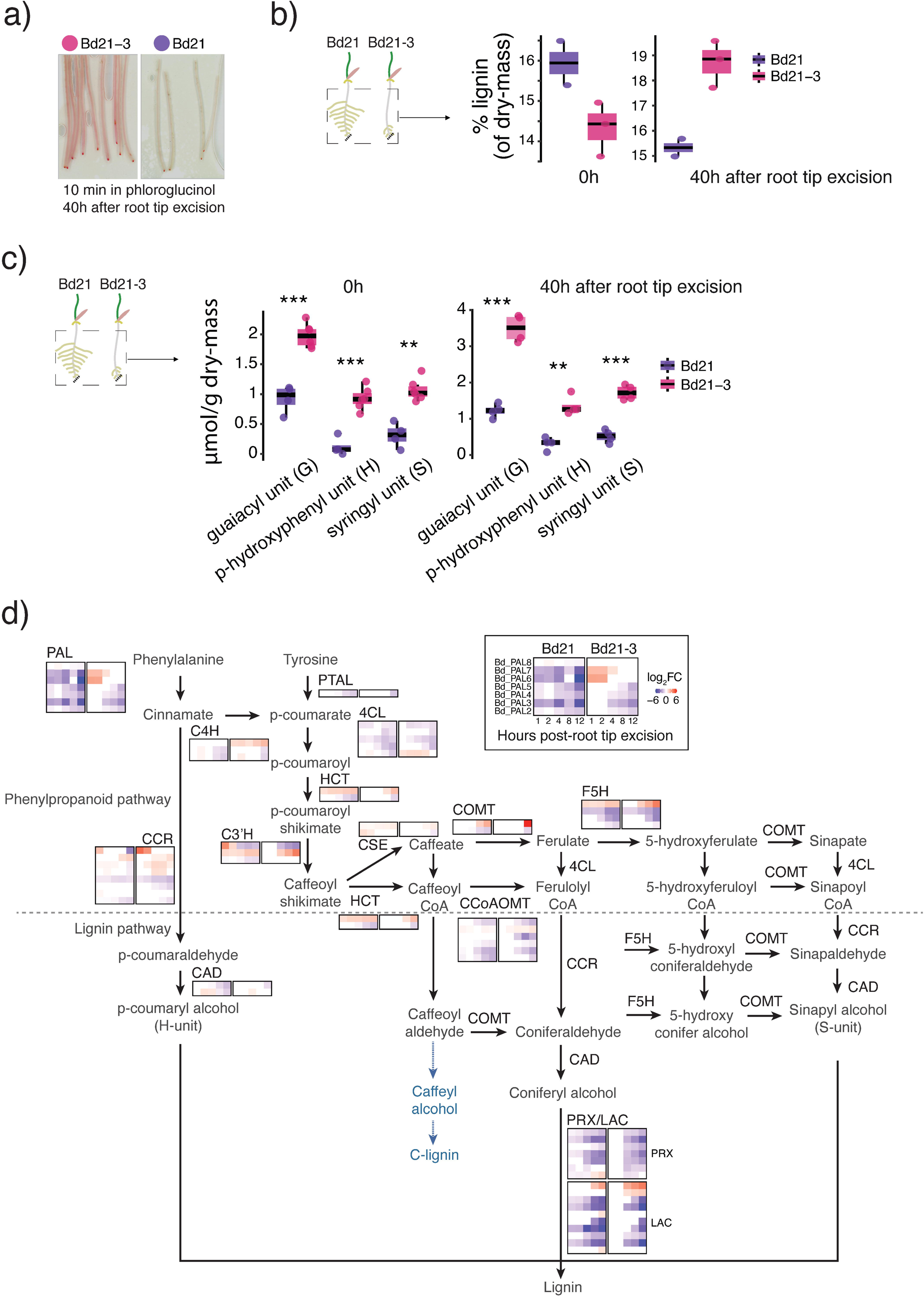
Bd21-3 “fishbone” root phenotype correlates with increased lignin deposition and upregulated lignin biosynthesis following root tip excision. (a) Phloroglucinol-HCl staining of roots at 40 hours after root tip excision reveals enhanced lignin accumulation in Bd21-3 compared to Bd21. Images are representative pooled individuals of three biological replicates. (b) Quantification of total lignin content by CASA in whole root systems at 0 h and 40 h after root tip excision. (c) Thioacidolysis-based quantification of lignin monomer composition (μmol/g dry weight) in whole roots at 0h and 40h after root tip excision, showing levels of guaiacyl (G), p-hydroxyphenyl (H), and syringyl (S) units. For each condition, roots from 30–50 seedlings were pooled per biological replicate (n = 3). In (b) and (c), each point represents a biological replicate. Statistical comparisons between genotypes at each time point were performed using two-sided Wilcoxon rank-sum tests (*p < 0.05; **p < 0.01; ***p < 0.001). (d) Schematic overview of the phenylpropanoid and lignin biosynthesis pathways. Heatmaps show logLJ fold changes in the expression of key biosynthetic genes at 1, 2, 4, 8, and 12 hours after root tip excision in Bd21 and Bd21-3.

To dissect the specific changes in lignin structure and content, we quantified monolignols using thioacidolysis. This revealed that Bd21-3 accumulated significantly higher levels of beta-O-4 linked H, G, and S units than Bd21, with a marked increase at 40 hours after root tip excision. We find a more pronounced increase in monolignols in the upper part of the primary root after root tip post excision of Bd21-3 plants where the lateral root emergence is delayed (Figs. 3c, S11). Together, the chemical analyses confirm that in Bd21-3, the “fishbone” phenotype is associated with a time-dependent and spatially defined increase in lignification. Focusing on the lignin biosynthetic pathway, we found that Bd21 and Bd21-3 differ in their transcriptomic response to lateral root synchronization (Fig. 3d). In Bd21-3, *BdPHENYLALANINE AMMONIA-LYASE 6* (*BdPAL6*) and *BdPAL7*, which encode enzymes catalysing the deamination of phenylalanine in the first step of the phenylpropanoid pathway, were both upregulated after root tip excision. This was accompanied by the upregulation of *BdCINNAMATE-4-HYDROXYLASE 3* (*BdC4H3*), indicating sustained flux through the lignin pathway. Further downstream, key genes involved in monolignol biosynthesis, including *Bd4-COUMARATE-COA LIGASE 1* (*Bd4CL1*) and *BdCINNAMOYL-COA REDUCTASES 1-4* (*BdCCR1–4*), showed higher fold-change in Bd21-3. Notably, induction of *BdCAFFEIC ACID O-METHYLTRANSFERASE 3* (*BdCOMT3*) and *BdFERULATE 5-HYDROXYLASE 5* (*BdF5H5*) suggested further progression toward monolignol production. Analysis of MYB transcription factors, which act as upstream regulators of secondary cell wall biosynthesis, revealed that *BdMYB108* and *BdMYB112* homologs were consistently upregulated in Bd21-3, while *BdMYB54* and *BdMYB74* were upregulated and *BdMYB5* and *BdMYB50* downregulated in Bd21 (Fig. S12). Altogether, these results indicate stronger transcriptional activation of lignin biosynthesis in Bd21-3 in response to root tip excision.

We then used Basic Fuchsin staining and 2-photon microscopy to achieve cellular resolution of the differential lignification. Confocal images pointed to the exodermis as the site of differential lignification between Bd21 and Bd21-3 (Fig. 4a). In Bd21-3, exodermal lignification was initiated earlier during lateral root development and more intensely compared to Bd21. By stage IV of lateral root development, the exodermis surrounding the emergence zone in Bd21-3 appeared already fully lignified, creating a continuous barrier. This premature reinforcement correlated with a flattened morphology of the lateral root primordia, suggesting that lignification promotes mechanical resistance (Fig. 4a). In contrast, exodermal lignification in Bd21 was delayed, allowing the lateral root to emerge with a typical elongated shape. Altogether, the lignin quantification and the 2-photon microscopy show that enhanced lignification in Bd21-3 is spatially targeted to the exodermis and occurs earlier during lateral root development, potentially forming a physical barrier that impairs emergence.

**Fig. 4.**
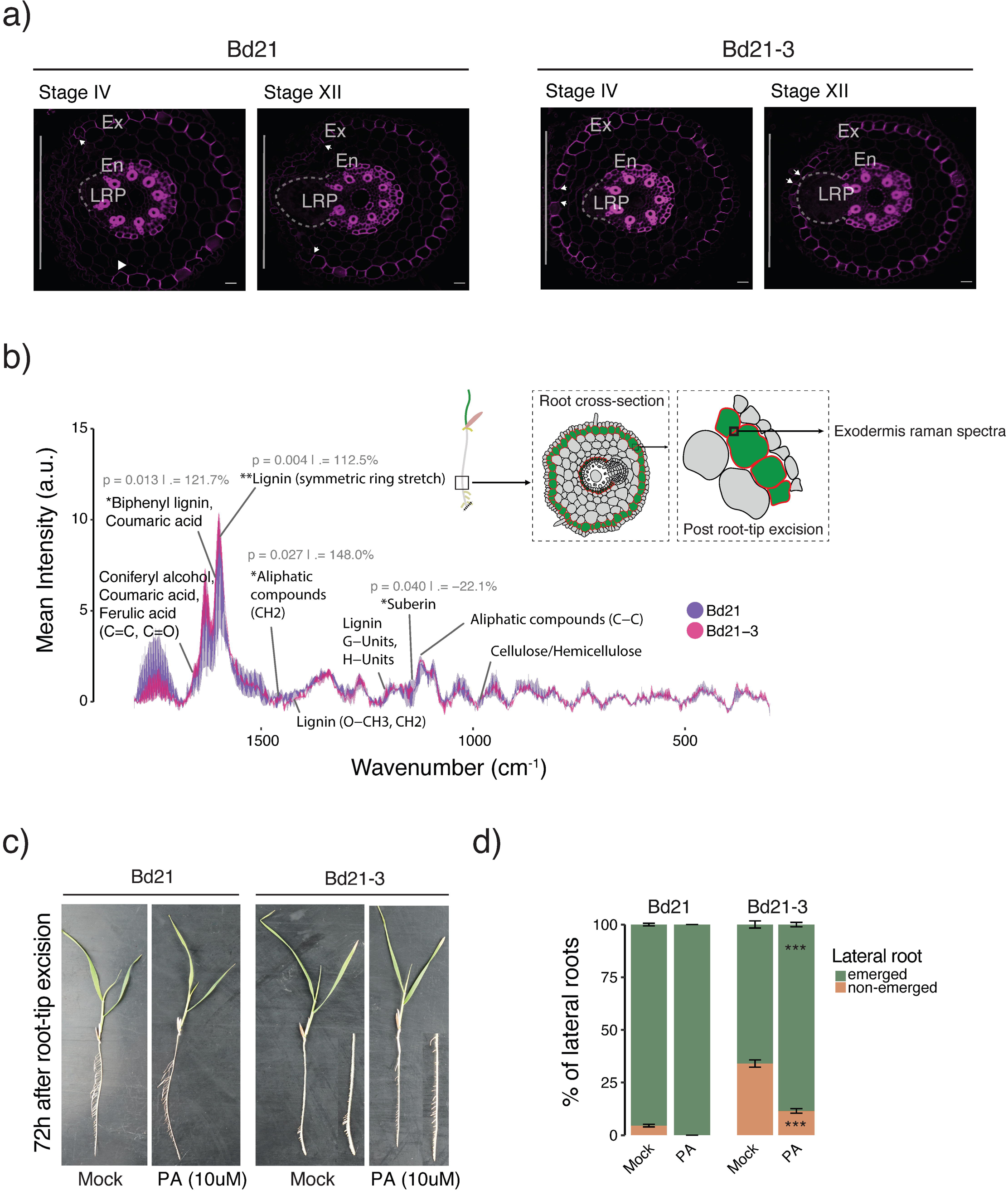
Bd21-3 exhibits enhanced and premature lignification of the exodermis during lateral root development. (a) Two-photon excitation microscopy images of Basic Fuchsin-stained root cross-sections from Bd21 (left panel) and Bd21-3 (right panel). A white vertical bar on the left side of each image marks the side facing the ½ MS medium. White arrowheads indicate the limits of exodermal lignification opposite the lateral root emergence side. Scale bars represent 50 μm. Forty-five seedlings were analysed, with three independent biological replicates for each staining approach. Ex: exodermis; En: endodermis; LRP: lateral root primordium. Scale bars: 20 µm. (b) Raman spectroscopy of the anticlinal cell walls of the exodermis region showing mean spectral intensities of major cell wall components in Bd21 and Bd21-3*. p*-values and relative intensity changes are indicated. (c) Representative images of whole root systems of Bd21 and Bd21-3 seedlings 72 hours after root tip excision treated with either mock or 10 µM piperonylic acid. (d) Distribution of emerged and non-emerged lateral roots under mock and piperonylic acid (PA) treatments. Bars show mean percentages (5–8 seedlings per condition, three biological replicates). Asterisks indicate significant differences between mock and piperonylic acid (PA) treatment, assessed using a binomial generalized linear mixed model with seedling as the unit and Petri dish as a random effect. Error bars represent standard error of the mean.

To gain precise *in situ* chemical insights, we conducted confocal Raman spectroscopy on the anticlinal cell walls of exodermal cells, which exhibited the strongest Basic Fuchsin staining (Fig. 4b). The resulting spectra revealed a profound chemical divergence between the two accessions. The exodermis of Bd21-3 exhibited significantly higher intensities for multiple peaks associated with lignin and phenolic compounds compared to Bd21. The most prominent lignin peak at ∼1600 cmLJ¹ (symmetric ring stretch), a proxy for total lignin content, was 112.5% higher in Bd21-3 (*p*=0.004). Furthermore, the peak at ∼1630 cmLJ¹, assigned to biphenyl lignin and coumaric acid, was 121.7% higher (*p*=0.013). In contrast, a peak associated with suberin at ∼1125 cmLJ¹ was significantly reduced by 22.1% in Bd21-3 (*p*=0.040). This shift in chemical profile points to a divergent strategy in outer tissue reinforcement between accessions.

To functionally confirm that increased lignification participates in inhibiting lateral root emergence in Bd21-3, we applied piperonylic acid (PA), a chemical inhibitor of the lignin biosynthesis enzyme C4H (Lee *et al*., 2013). In cleared roots, mock-treated Bd21-3 plants displayed a significantly higher proportion of non-emerged lateral roots compared to the wild-type Bd21. Treatment with PA (10 µM) markedly reduced this proportion in Bd21-3, restoring emergence levels closer to those observed in Bd21 (Figs. 4c, d). By contrast, Bd21 plants maintained high levels of emergence under both mock and PA conditions. This corresponds to a partial, though incomplete, shift from the inhibited “fishbone” root system architecture toward a more developed “pine tree” phenotype, demonstrating that inhibition of lignin biosynthesis partially rescues the defective emergence in Bd21-3. Collectively, our results indicate that the Bd21-3 accession responds to root tip excision with a specialized program of rapid and intense lignification in the exodermis, likely creating a rigid physical barrier that restricts lateral root emergence and alters overall root system architecture (Fig. 5).

**Fig. 5.**
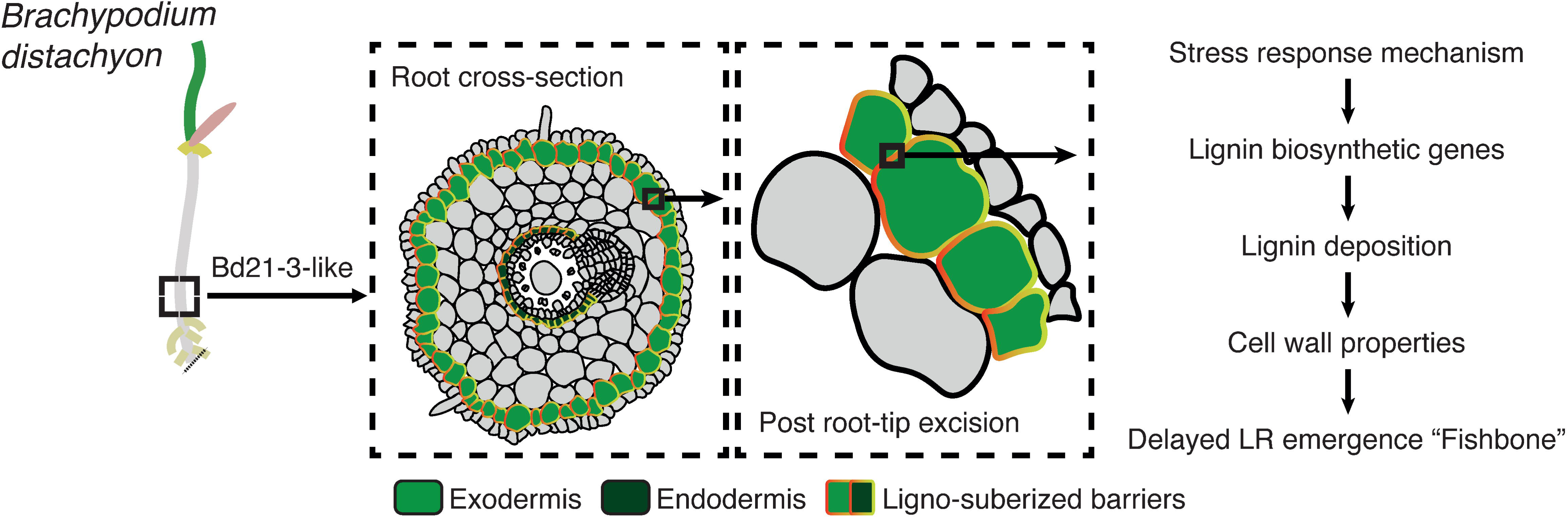
Schematic model for the stress-induced inhibition of lateral root emergence in *B. distachyon* accession Bd21-3. This model illustrates the proposed mechanism underlying the “fishbone” phenotype. A stress event, such as root tip excision, initiates a specific response pathway in the outer root tissues, particularly the exodermis. This response triggers the transcriptional activation of lignin biosynthetic genes, leading to increased lignin deposition within the exodermal cell walls. The resulting change in cell wall properties increases their rigidity, creating a mechanical constraint that ultimately causes delayed lateral root emergence.

## Discussion

Here, we uncovered striking natural variation in RSA plasticity in response to root tip excision between two *B. distachyon* accessions. We highlight two contrasting architectures, with Bd21 displaying a “pine tree” phenotype marked by evenly spaced lateral roots along the primary root, while Bd21-3 exhibits a “fishbone” phenotype in which lateral roots emerge proximally but are delayed in the upper part of the primary root after root tip excision. Importantly, we show that the phenotypic divergence is not due to defects in lateral root initiations but rather to a block in emergence, specifically in the upper part of the primary root zone (Fig. 1).

Root tip excision has long been known to stimulate lateral roots formation and emergence across various plant species (Torrey, 1950; Thomas *et al*., 2014; Kawai *et al*., 2017). In some case, the differential emergence of lateral roots in proximal and upper regions may reflect differences in the developmental stage of lateral roots along the root axis. In rice for instance, seminal root tip excision induces expansion of lateral root primordia through enhanced periclinal divisions in the ground tissue, but also a broader stele and this response is most pronounced in lateral root primordia closest to the excision site (Kawai *et al*., 2017). Furthermore, in rice, lateral roots identity is defined by two distinct subtypes: long, highly branched L-type lateral roots and short, unbranched S-type lateral roots. Experiments involving root tip excision have shown that antagonistic interactions of *WUSCHEL-RELATED HOMEOBOX* (*WOX*) transcription factors control the balance between L-type and S-type lateral roots, promoting the formation of L-type lateral roots while suppressing S-type lateral roots development (Kawai *et al*., 2022). It would be valuable to explore whether *B. distachyon* also produces different lateral root types *e.g.* L-type and S-type and how their distribution varies along the primary root in relation to proximity to the excision site. The morphology and type of lateral roots could directly influence both the growth-rate through overlying tissues and the timing of emergence. Thus, the delay in lateral roots emergence associated with the Bd21-3 “fishbone” phenotype may reflect not only changes in cell wall *i.e.* mechanical constraints but also differences in lateral root primordia identity and development.

Our results suggest that divergence in RSA could also arise, at least in part, from intraspecific genomic variation segregating among *B. distachyon* accessions, potentially contributing to local adaptation. Notably, differences in the expression of single-copy orthologous genes were observed during the later stages of the root tip excision response (Fig. 2a). Furthermore, GO enrichment analysis of genes within multicopy orthogroups specific to the Bd21 accession revealed a strong overrepresentation of biological processes related to DNA repair and cellular responses to stress (Fig. S4). Bd21 may possess an expansion of stress-responsive gene families that enable more effective mitigation of cellular damage caused by wounding, facilitating a more developmentally permissive root program. In *B. distachyon*, a pangenome built from 54 accessions showed that approximately 65% of the gene pool is accessory, with many genes implicated in defence and development-related functions (Gordon *et al*., 2017). This same pangenome revealed that gene presence/absence and copy number variation across accessions are tightly linked to population structure and adaptive traits. Further functional and population-level studies would help determine whether the two phenotypes are linked to adaptive strategies shaped by natural selection acting on the species’ pangenome.

Our results highlighted pathway-wide reprogramming towards intensive lignin biosynthesis, both transcriptionally and biochemically. We find that following root tip excision, the root exodermis in the Bd21-3 accession accumulated significantly more lignin characterized by biphenyl 5−5’ cross-links derived primarily from guaiacyl (G) units (Boerjan *et al*., 2003; Guadix-Montero & Sankar, 2018). The prevalence of G-units is known to promote a more cross-linked and condensed polymer structure, resulting in a stiffer, more recalcitrant matrix that can function as an effective physical barrier (Fig. 4b) (Rosado *et al*., 2021; Balk *et al*., 2023; Muretta *et al*., 2024). Furthermore, under stress-like conditions, H-unit–rich lignin has been shown to accumulate (Tobimatsu & Schuetz, 2019; Gladala-Kostarz *et al*., 2020). The increase in H-monomers observed in Bd21-3 likely reflects a similar cell wall stress response, triggered here by root tip excision. Although H-units are less commonly incorporated in developmental lignin, their presence is frequently associated with increased polymer rigidity and resistance to enzymatic degradation (Mamedes-Rodrigues *et al*., 2019; Balk *et al*., 2023). Notably, mechanical constraints resulting from this altered lignin composition has been linked to increased tissue brittleness, independent of total lignin quantity (Timpano *et al*., 2015). These findings show that the spatial distribution and chemical composition of lignin influence the mechanical behaviour and developmental outcomes.

In our study, these structural consequences are most apparent at the exodermis, where Bd21-3 showed early and intense exodermal lignification directly above developing lateral root, forming a continuous barrier (Fig. 4a). This lignified layer coincided with flattened, stalled primordia and a significant reduction in successful emergence events, *in fine* resulting morphologically as the “fishbone” phenotype. Conversely, Bd21 displayed a more permissive response: exodermal lignification was delayed and localized, creating temporary openings that allowed lateral roots to emerge normally. Furthermore, the functional relevance of this mechanical barrier was further validated by pharmacological inhibition of lignin biosynthesis using piperonylic acid, which partially restored lateral roots emergence and root architecture in Bd21-3 (Fig. 4c, d). Together, our data supports a model in which Bd21-3 activates a lignin-based fortification program in response to mechanical stress, characterized by early and localized deposition of G- and H-rich lignin polymers in the exodermis. Notably, the concurrent increase in S-units, typically associated with developmental lignin or hydrophobic sealing functions, suggests that both stress and developmental pathways may be co-opted in Bd21-3, possibly resulting in a lignin barrier that is both hydrophobic and mechanically rigid (Vanholme *et al*., 2012; Barros *et al*., 2015). The reinforcement of the outer root tissues in Bd21-3 likely impacts the flexibility needed to accommodate lateral root emergence. In *A. thaliana*, compromised Casparian strip integrity and low calcium availability promotes ectopic lignin deposition in the endodermis in a SGN3/GSO1-dependent manner (Lai *et al*., 2025). This excessive barrier lignification mechanically impedes lateral root emergence, defining a developmental checkpoint in which lignin functions as a mechanical gatekeeper (Banda *et al*., 2019). In Bd21-3, premature and spatially excessive lignification of the exodermis is strongly induced as a protective response to wounding and similarly acts as a mechanical constraint on organ emergence. Whether a surveillance system is actively involved in the balance between organogenesis and the response to wounding in both accessions remains unknown.

Other plant species have shown similar plasticity of their root exodermis cell layer. In tomato (*Solanum lycopersicum*), the exodermis does not form a Casparian strip but instead develops a structurally distinct polar lignin cap that functions as a selective apoplastic barrier (Manzano *et al*., 2024). Although functionally analogous to the endodermal Casparian strip, the regulation of this polar lignin cap involves a genetically distinct module, and its formation is spatially restricted to the outermost cortical layer by the repressive action of the transcription factors *SlSCHIZORIZA* (*SlSCZ*) and *SlEXO1*. Observations on different plant species therefore suggest a form of functional convergence, where lignin-based barriers with comparable physiological roles can emerge through unique regulatory pathways. In addition to the lignified barrier, exodermis suberization is also critical for drought tolerance in tomato, operating as a functional equivalent to the *A. thaliana* endodermis (Manzano *et al*., 2024). In tomato, the conserved suberin biosynthetic pathway, including regulators like *SlMYB92*, has been evolutionarily rewired to act in the exodermis rather than the endodermis (Cantó-Pastor *et al*., 2024). Recently, ABA was shown to first activate general stress-responsive elements and then induce lignin and suberin biosynthesis through MYB transcription factors in chickpea (Jo *et al*., 2025). In our dataset, several MYB genes known to be ABA-responsive, such as *MYB94*, *MYB91*, *MYB108*, and *MYB112*, were constitutively upregulated in Bd21-3, while others like *MYB93*, a negative regulator of lateral root emergence (Xiao *et al*., 2021; Uemura *et al*., 2023), were downregulated. These patterns would point to ABA as a key upstream regulator of the altered MYB landscape in Bd21-3.

In *A. thaliana*, cell wall remodelling plays a central role in lateral root emergence (Swarup *et al*., 2008; Péret *et al*., 2012; Kumpf *et al*., 2013; Lucas *et al*., 2013; Lewis *et al*., 2013; Vermeer *et al*., 2014; Roycewicz & Malamy, 2014; Berhin *et al*., 2019; Wachsman *et al*., 2020; Ursache *et al*., 2021). Consistent with this, our time-course transcriptomic analysis revealed dynamic regulation of numerous cell wall modifying enzyme families, particularly glycoside hydrolases, in both *B. distachyon* accessions after root tip excision (Fig. 2d–f; Fig. S7–S8). This suggests a tightly orchestrated remodelling process, potentially involving both cell wall loosening and reinforcement in a spatially and temporally regulated manner across different tissue layers. Recent studies have underscored that lateral root emergence depends on precise and layer-specific remodelling of the cell wall (Wachsman *et al*., 2020; Ursache *et al*., 2021). A key future challenge will be to integrate single-cell transcriptomics with high-resolution cell wall chemical profiling, enabling the mapping of transcriptional programs directly to physical changes in tissue properties. Moreover, genome-wide association studies (GWAS) across diverse *B. distachyon* accessions could help pinpoint the genetic loci responsible for variation in the root system architecture. In conclusion, our study reveals that, in response to wounding, exodermal lignification acts as a dynamically regulated mechanical barrier that can restrict lateral root emergence, driving intraspecific variation in the overall RSA. Far from being a passive structural layer, the exodermis emerges as a key regulatory interface, responsive to both intrinsic developmental signals and extrinsic stress cues. These findings not only advance our understanding of root development in grasses but also open promising avenues for engineering stress-resilient crops by targeting the regulatory pathways that govern root barrier plasticity.

## Acknowledgements

We would like to thank the Botanical Garden of Neuchâtel and the University of Neuchâtel for their support in propagating and maintaining the *B. distachyon* accessions collection. K.B. was supported by a Marie Skłodowska-Curie Global Fellowship (PLANTiD, Grant Agreement No. 101106663). Work in the Vermeer lab was funded by the Swiss National Science Foundation (project numbers 157524 and 197568 awarded to J.E.M.V.) and by the University of Neuchâtel.

## Author contributions

CDJVT and KB characterised the root system architecture phenotypes in *B. distachyon* accessions. AR provided the Mediterranean *B. distachyon* accessions. CDJVT and KB generated the seed resources for these accessions, with support from the Botanical Garden of Neuchâtel. CDJVT prepared the RNA samples with assistance from KB. TB performed the genomic analyses and processed the transcriptomic data. LP, RS, and KB conducted lignin quantification. AO and KB carried out the Raman spectroscopy analysis. CDJVT and KB performed the brightfield and confocal imaging. KB, TB, and JEMV conceived and designed the research. TB prepared the figures with input from KB. KB and TB co-wrote the first draft of the manuscript. KB, TB, and JEMV supervised the project.

## Data availability

The data that support this study are available within the article as part of the Supplementary Tables. The RNA-sequencing raw sequencing data was deposited at the NCBI Short Read Archive under the accession number PRJNA1312106.

## Supplementary tables

**Table S1: Root phenotyping data for *B. distachyon* accessions**. This table contains the related “fishbone” or “pine tree” phenotypes for each of the 23 *B. distachyon* accessions. Data was collected from seedlings grown on agar plates, as described in the Materials and Methods. Root systems were scanned and the phenotype analyzed. Related to Fig. 1b.

**Table S2: Root length data for *B. distachyon* accessions**. The table contains the raw measurements of primary root length, numbers of emerged lateral roots and lateral root density for the Bd21 and Bd21-3 *B. distachyon* accessions. Data was collected from seedlings grown on agar plates, as described in the Materials and Methods. Root systems were scanned and root lengths analyzed using ImageJ. Related to Fig. 1c.

**Table S3: Number of emerged lateral roots after root tip excision**. The table provides the lateral root number measured for specific 1-cm regions along the primary root (0-1 cm, 1-2 cm, etc.) for accessions Bd21 and Bd21-3, after root tip excision. Data were collected from seedlings grown on agar plates, cleared and phenotyped as described in the Materials and Methods. Related to Fig. 1d.

**Table S4: Number of emerged lateral roots after root-tip excision**. The table provides the lateral root number measured for specific 1-cm regions along the primary root (0-1 cm, 1-2 cm, etc.) for accessions Bd21 and Bd21-3, after root tip excision. Data were collected from seedlings grown on agar plates, cleared and phenotyped as described in the Materials and Methods. Related to Fig. 1d.

**Table S5: Proportions of lateral root primordia stages in the primary root at in different zones after root tip excision**. Raw data for Bd21 and Bd21-3 *B. distachyon* accessions. Related to Fig. S2. This table provides the proportions of lateral root primordia stages observed in specific zones of the primary root after root tip excision for the Bd21 and Bd21-3 *B. distachyon* accessions. Data were collected from seedlings grown on agar plates, cleared and phenotyped as described in the Materials and Methods.

**Table S6: Orthology assignment between Bd21 and Bd21-3 proteins as per orthofinder output**. The “n_isoforms” column denotes the number of protein isoforms annotated for that gene. Finally, “pancategory” classifies the orthogroup based on copy number across genomes, where “single-copy” refers to one-to-one orthologs present in all genomes, and “multi-copy” indicates the presence of multiple paralogs in at least one genome.

**Table S7: Summary table of the differential gene expression analysis performed using DESeq2 across multiple time points following root tip excision, relative to the baseline (0 hours)**. The “condition” column indicates the time point in the kinetic series (1, 2, 4, 8, 12 hours) compared to 0 hours. “pancategory” refers to the pangenome category of the gene, describing whether it is a single-copy ortholog or present in multiple accessions, and “deg” indicates whether the gene is significantly differentially expressed at a 5% false discovery rate.

**Table S8: Comparison of differential gene expression for single-copy orthologs between the two *B. distachyon* accessions across multiple time points following root tip excision**. Each row corresponds to a specific orthogroup at a given time point (“condition”). Columns “Bd21” and “Bd21_3” report the log2 fold-change values of the corresponding gene(s) in each accession, with missing values (NA) indicating that no significant differential expression was detected in that accession. The “category” column specifies whether the gene is significantly differentially expressed in one or both accessions.

**Table S9: Summary of word occurrences in the Pfam description of the top 100 up- and down-regulated genes in the full transcriptomic dataset.**

**Table S10: Gene ontology enrichment at the biological process level of genes similarly differentially expressed in both accessions**. Fisher’s exact test was performed using topGO in R for each time-point (condition).

**Table S11: Differential expression of a selection of cell-wall related enzyme families and their functional annotation in the WallProt database.**

**Table S12: Gene ontology enrichment at the molecular function level of differentially expressed genes**. Fisher’s exact test was performed using topGO in R for genes differentially expressed in both accessions (co_deg), Bd21 only (Bd21_deg), Bd21-3 only (Bd21_3_deg) or in opposite direction (op_deg).

**Table S13: Differential expression of cell wall–related gene families**. The table shows the differential expression profiles of cell wall–related gene families following root tip excision in Bd21 and Bd21-3. LogLJ fold-change expression values are presented across a 12-hour time course in Bd21 and Bd21-3. Each subplot corresponds to one gene family, with individual gene trajectories displayed as lines. Pie charts above each family indicate the proportion of genes significantly upregulated or downregulated. Related to Fig. S8.

**Table S14: Quantification of total lignin content by CASA in whole root systems at 0 h and 40 h after root tip excision**. The table provides the percentage of lignin per dry weight in the primary root after root tip excision for the Bd21 and Bd21-3 *B. distachyon* accessions. Data were collected from seedlings grown on agar plates; root were harvested as described in the Materials and Methods. Related to Fig. 3b.

**Table S15: Quantification of total lignin content by thioacidolysis in the primary roots at 0 h and 40 h after root tip excision**. The table provides the quantity of lignin per μmol/g dry-mass in the primary root after root tip excision for the Bd21 and Bd21-3 *B. distachyon* accessions. It includes measurements of lignin content and the molar ratio of S-lignin and G-lignin monomers. Data were collected from seedlings grown on agar plates; root were harvested as described in the Materials and Methods. Related to Fig. 3c.

**Table S16: Differential expression of lignin biosynthesis-related genes**. The table provides log_2_ fold-change values for the subsets of genes involved in the phenylpropanoid and lignin biosynthesis pathway at 1, 2, 4, 8, and 12 hours after root tip excision. Related to Fig. 3d.

**Table S17: Quantification of total lignin content by thioacidolysis in specific zones of the primary roots at 0 h and 40 h after root tip excision**. The table provides the quantity of lignin per μmol/g dry-mass in the primary root after root tip excision for the Bd21 and Bd21-3 *B. distachyon* accessions. It includes measurements of lignin content and the molar ratio of S-lignin and G-lignin monomers. Data were collected from seedlings grown on agar plates; root were harvested as described in the Materials and Methods. Related to Fig. 3c.

**Table S18: Differential expression of MYB-related transcriptional factor genes**. The table provides log_2_ fold-change values for a subset of genes annotated as part of the MYB transcription factor family. Related to Fig. S12.

**Table S19: Summary statistics of the RAMAN spectra at the cell-walls of the root exodermis.**

**Table S20: Proportions of emerged versus non-emerged lateral root for Bd21 and Bd21-3 seedlings treated with either mock solution or 10 µM piperonylic acid (PA).** The table provides the number of emerged and non-emerged lateral root primordia measured in the whole primary root after root tip excision for the Bd21 and Bd21-3 *B. distachyon* accessions treated with either mock solution or 10 µM piperonylic acid (PA). Data were collected from seedlings grown on agar plates, cleared and phenotyped as described in the Materials and Methods. Related to Fig. 4d.

