## Supplementary Tables for "Exodermis lignification impacts lateral root emergence in *Brachypodium distachyon*"

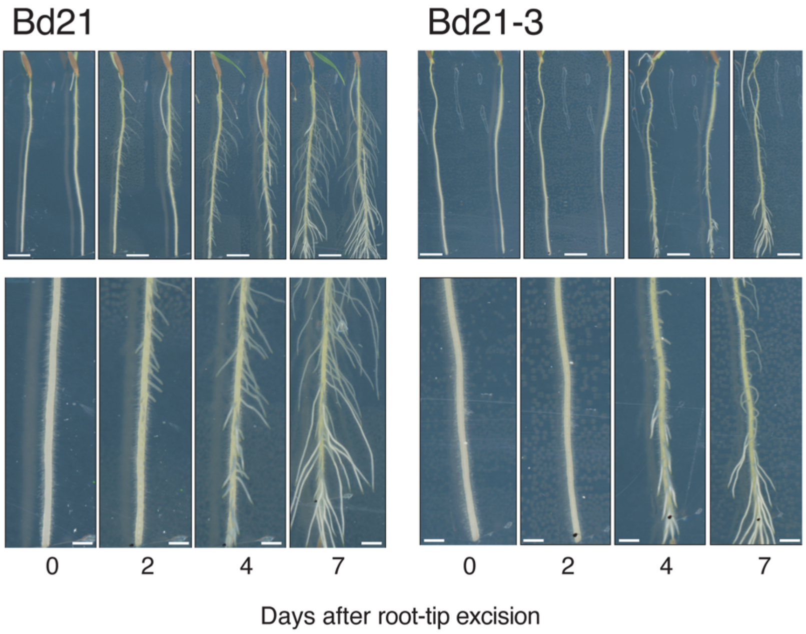


**Figure S1. Bd21 and Bd21-3 display distinct lateral root emergence dynamics following root tip excision.** Time-course of lateral root emergence dynamics after root tip excision. Representative images of whole root systems of Bd21 and Bd21-3 at 0, 2, 4, and 7 days after root tip excision. The lower panels provide magnified views showing the emergence of lateral roots along the entire root in Bd21 and the restricted emergence in the upper regions of Bd21-3. Scale bars: Upper panel, 5 mm; lower panel, 3 mm.

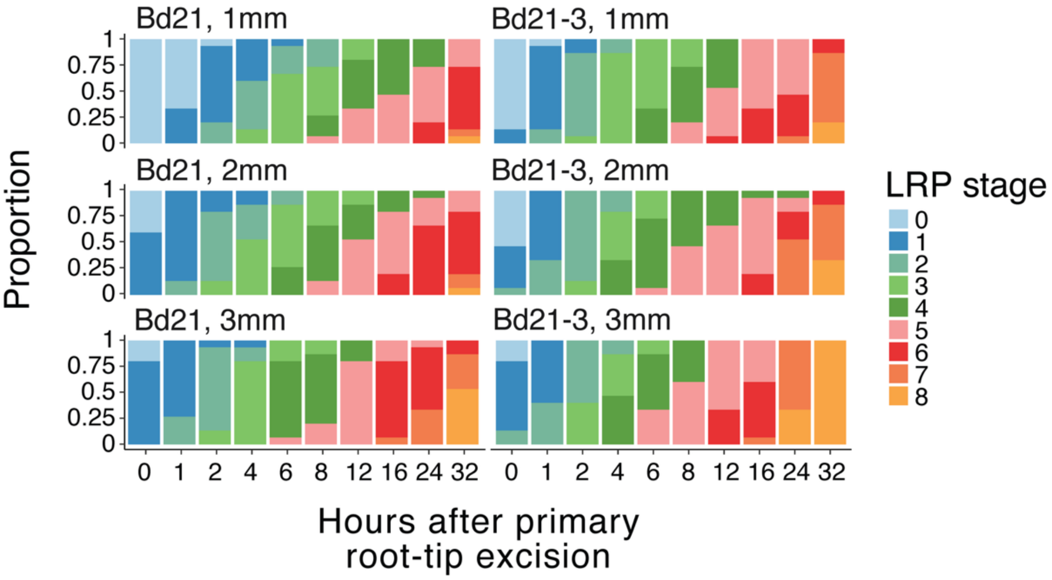


**Figure S2. Lateral root primordium development is synchronized and comparable between accessions.** Quantification of lateral root primordium stages within the first 1, 2, and 3 mm of the root above the excision site over a 32-hour time course. The stacked bar charts show the proportion of lateral root primordia at each developmental stage (0-8) for Bd21 and Bd21-3.

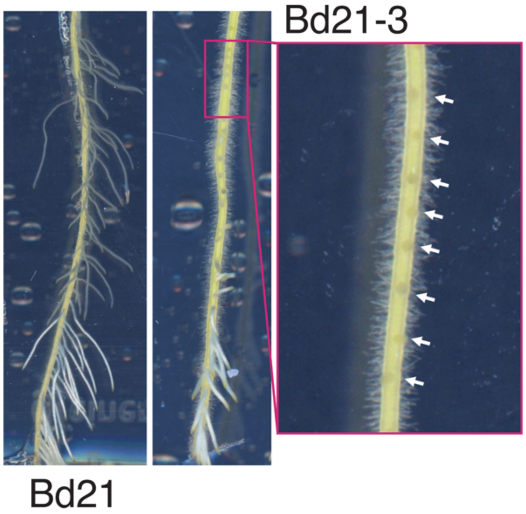


**Figure S3. Bd21 “pine-tree” and Bd21-3 “fishbone” root phenotypes 60h after root tip excision.** Magnified view of non-emerged lateral root primordia in Bd21-3. Representative images of Bd21 and Bd21-3 roots at 60 hours after root tip excision.

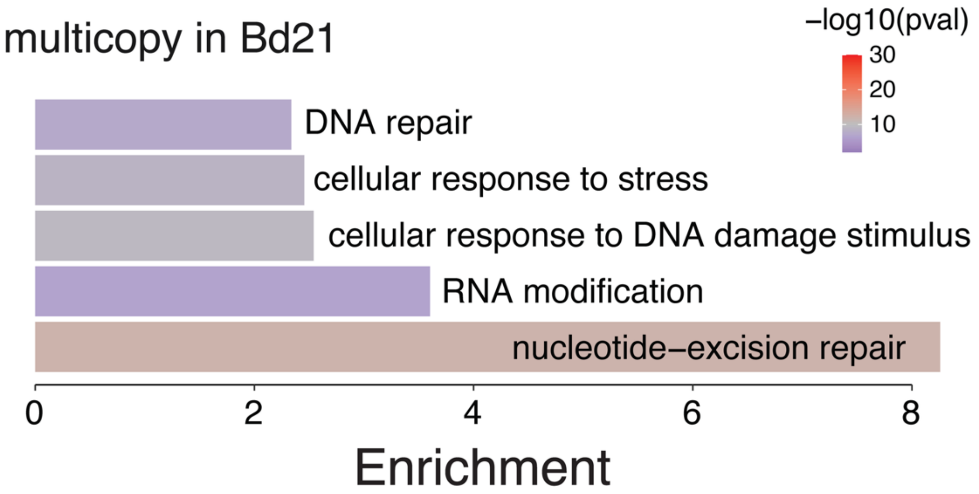


**Figure S4. Orthogroups with multiple copies in the Bd21 accession are enriched for stress and DNA repair functions.** GO enrichment analysis for biological processes associated with genes found in multicopy orthogroups specific to the Bd21 accession. The bar plot shows the enrichment scores for the most significantly overrepresented GO terms, which are predominantly related to DNA repair and cellular responses to stress. The colour of each bar corresponds to the statistical significance ($-\log_{10}$(p-value)).

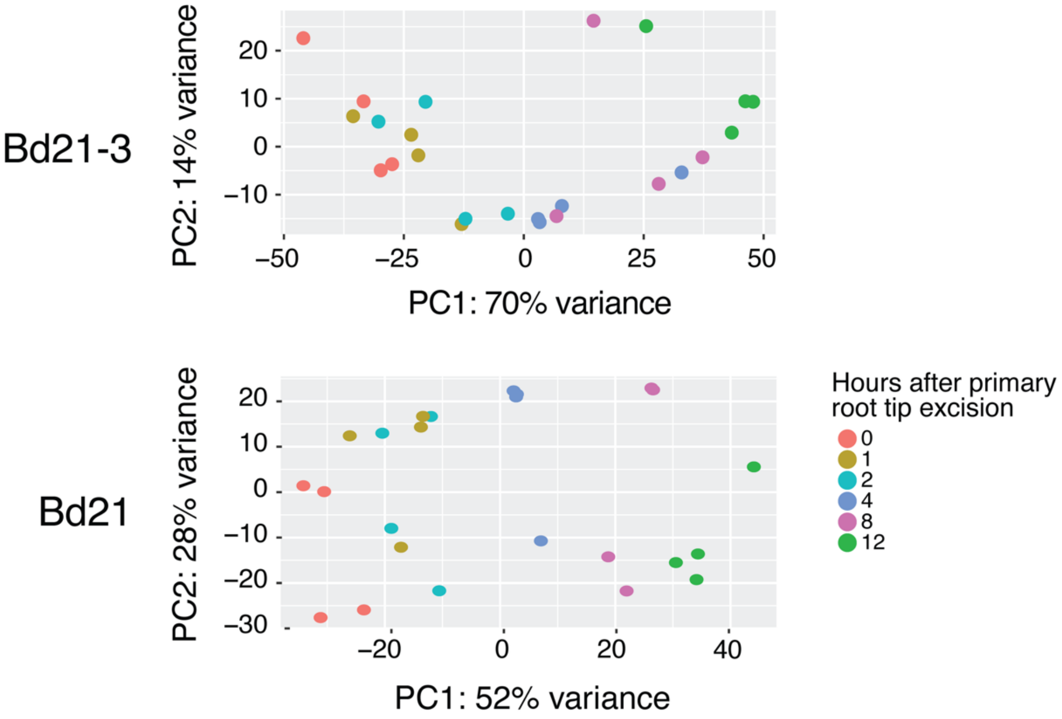


**Figure S5. RNA-seq analysis highlights distinct early and late transcriptional responses in both *B.distachyon* accessions.** Principal component analysis of RNA-seq data from root tissue collected at 0, 1, 2, 4, 8, and 12 hours post excision.

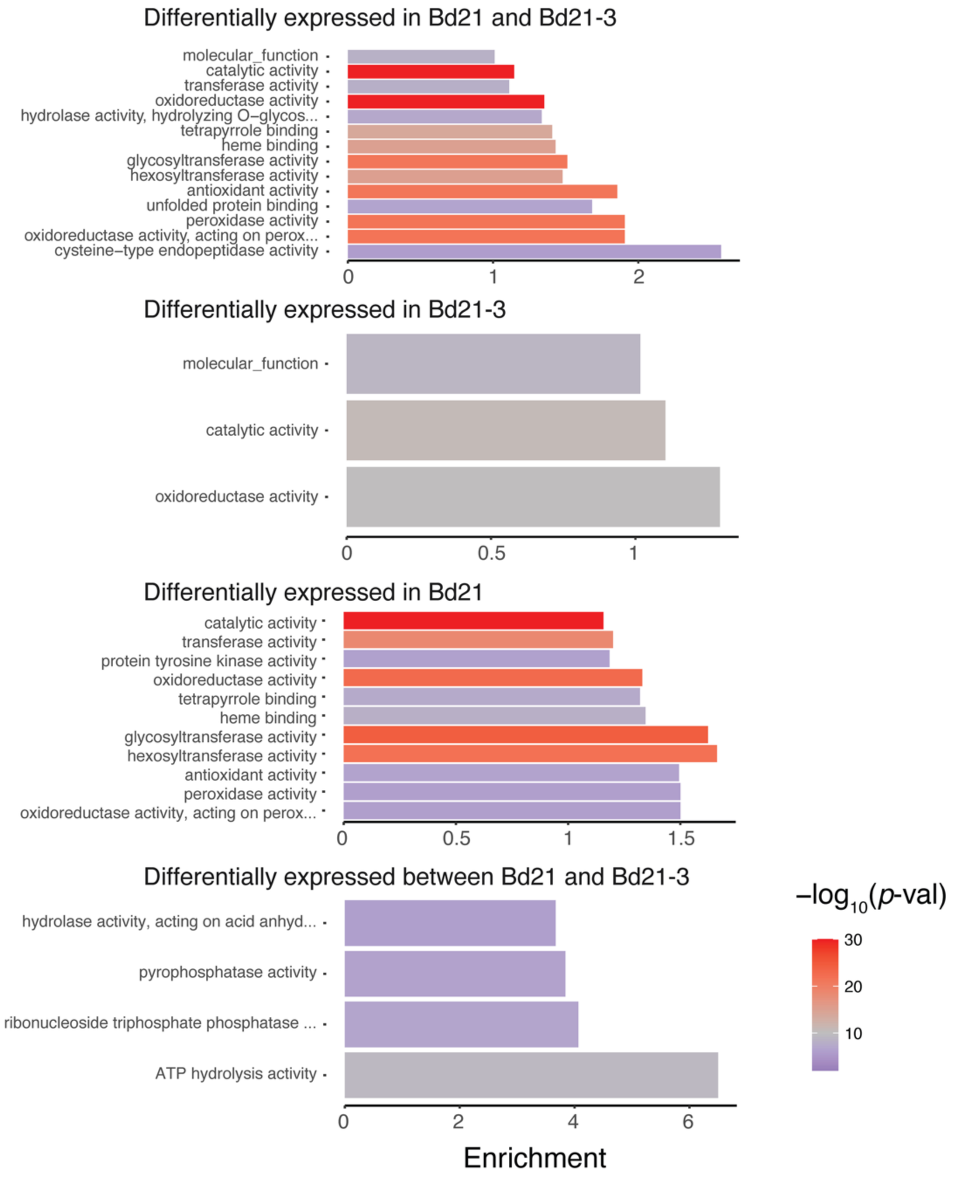


**Figure S6. Gene Ontology enrichment analysis for differentially expressed genes.** The bar charts display the GO enrichment analysis for different subsets of differentially expressed genes following root tip excision, focusing on the "molecular function" category. The x-axis represents the enrichment score, and the colour of the bars indicates the statistical significance (-log10(p-value)). (Top panel) GO terms enriched for genes that are differentially expressed in both *Bd21* and *Bd21-3*. (Second panel) GO terms enriched for genes differentially expressed only in *Bd21-3*. (Third panel) GO terms enriched for genes differentially expressed only in *Bd21*. (Bottom panel) GO terms enriched for the 240 genes that show opposite regulation between the two accessions.

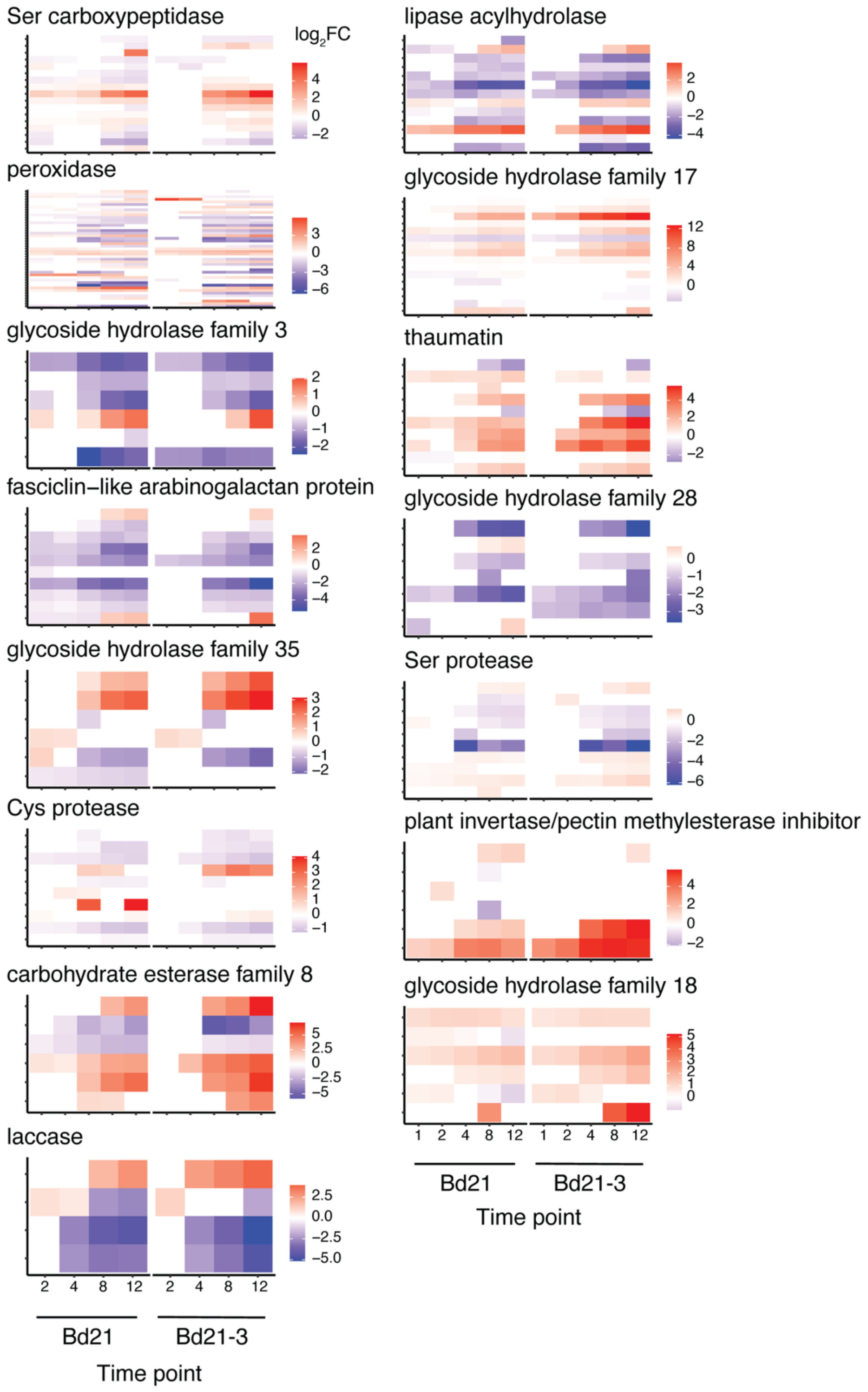


**Figure S7. Comparative transcriptomic analysis reveals a divergent cell wall remodelling response in Bd21 and Bd21-3 after root tip excision.** Heatmaps showing the expression dynamics ($\log_2$ fold-change) for individual genes within selected cell wall-related enzyme families. Each row represents a single gene, and its expression is shown over the 12-hour time course for both Bd21 and Bd21-3.

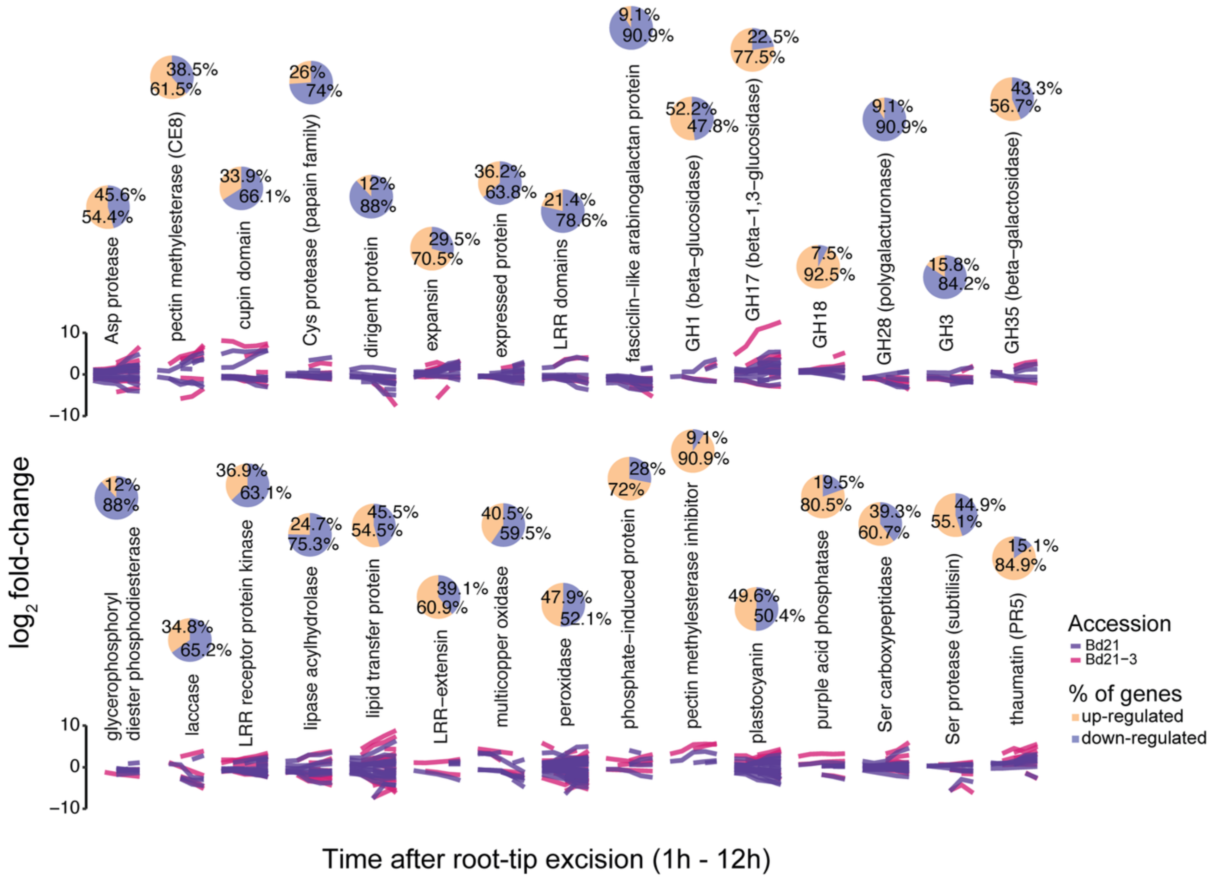


**Figure S8**. **Differential expression profiles of cell wall–related gene families following root tip excision in Bd21 and Bd21-3.** $Log_2$ fold-change expression profiles of genes grouped by annotated protein family across a 12-hour time course after root tip excision in Bd21 (purple) and Bd21-3 (magenta). Each subplot represents one gene family, with individual gene trajectories shown as lines. Pie charts above each family indicate the percentage of genes that are significantly upregulated (orange) or downregulated (blue).

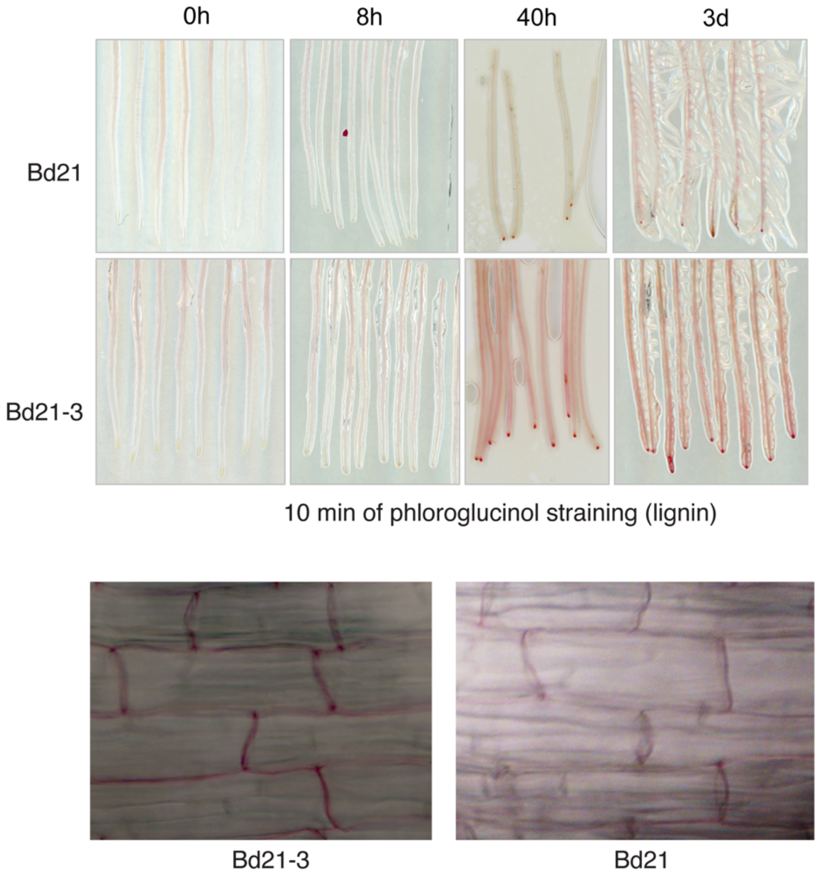


**Fig. S9. Spatio-temporal analysis of lignin deposition after root tip excision. This figure visualizes the dynamics of lignin deposition using phloroglucinol staining.** The top panel shows a time course at 0, 8, 40 hours, and 3 days after root tip excision in the roots of Bd21 and Bd21-3. The bottom panel provides magnified images of the root epidermis, highlighting the strong and distinct lignification of cell walls in Bd21-3 compared to the faint staining in Bd21.

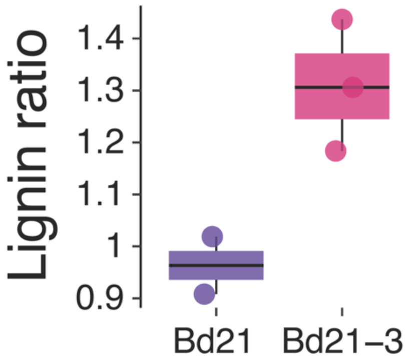


**Fig. S10. Baseline lignin content is comparable between Bd21 and Bd21-3.** Quantification of total lignin content (%) in whole roots of Bd21 and Bd21-3 as determined by the CASA method. Each point represents a biological replicate.

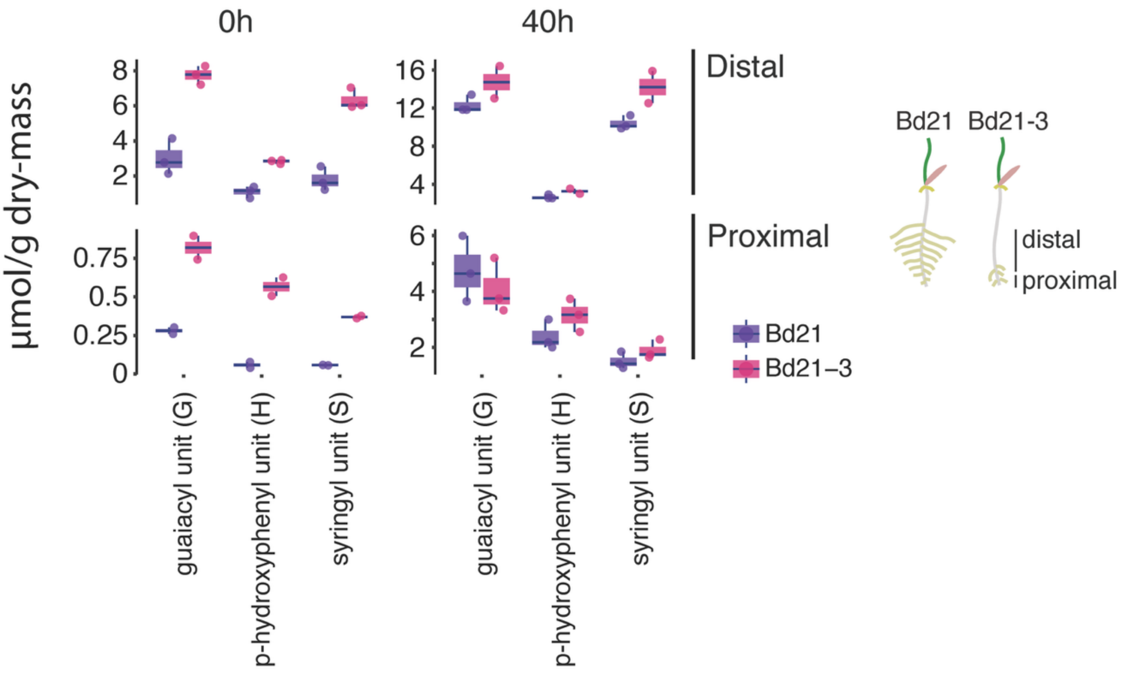


**Fig. S11**. **Lignin monomer composition is elevated in the upper root zone of Bd21-3.** Spatially resolved quantification of lignin monolignol composition in proximal and upper root regions at 0- and 40-hours post excision. The box plots show the amounts (µmol/g) of guaiacyl (G), p-hydroxyphenyl (H), and syringyl (S) units.

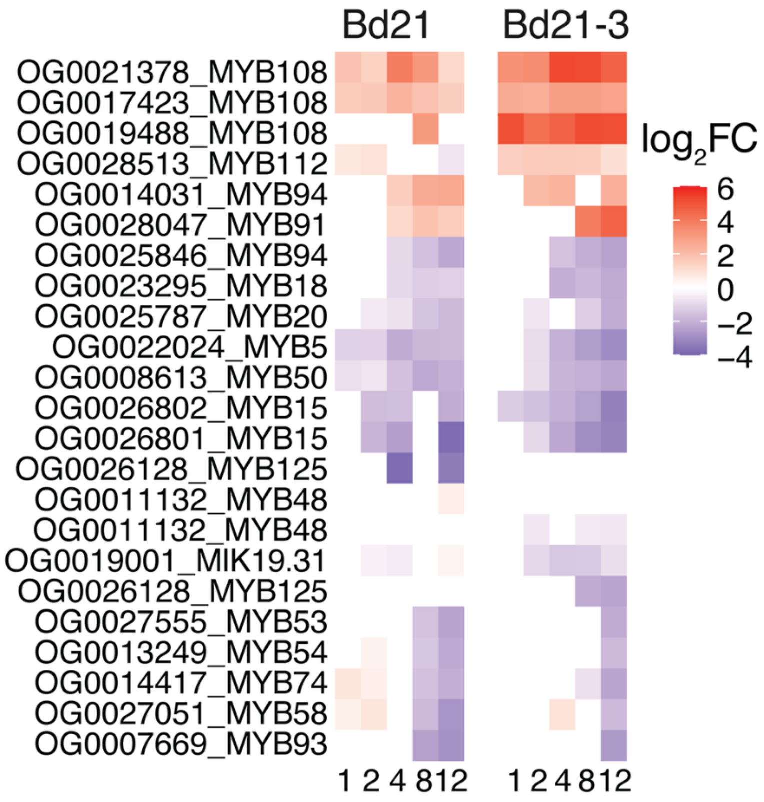


**Fig. S12. Expression dynamics of selected MYB transcription factor homologs following root tip excision**. The heatmaps display the $log_2$ fold-change at 1, 2, 4, 8, and 12 hours for Bd21 and Bd21-3.
